# Non-overlapping requirements of ASNA-1 function for insulin secretion, cisplatin resilience, and growth revealed by genetic analysis of point mutants in *C. elegans*

**DOI:** 10.1101/2022.07.12.499739

**Authors:** Dorota Raj, Agnieszka Podraza-Farhanieh, Pablo Gallego, Gautam Kao, Peter Naredi

## Abstract

ASNA1 plays an essential role in cisplatin chemotherapy response, type 2 diabetes, and heart disease. It is also an important biomarker in the treatment response of many diseases. Biochemically, ASNA1 has two mutually exclusive redox modulated roles: a tail-anchored protein (TAP) targeting function in the reduced state and a holdase/chaperone function in the oxidized state. Assigning biochemical roles of ASNA-1 to biomedical functions is crucial for successful therapy development. Our previous work showed the relevance of the *C. elegans* ASNA-1 homolog in modeling cisplatin response and insulin secretion. Here we analyzed two-point mutants in highly conserved residues in *C. elegans* ASNA-1 and identified their importance in separating cisplatin response from its roles in insulin secretion. Further, using targeted depletion we showed *asna-1* tissue requirements for *C. elegans* growth and development. We concluded that, targeting single residues in ASNA-1 affecting Switch I/II domain function, in comparison to complete knockdown counteracted cisplatin resistance without jeopardizing other important biological functions. Taken together, our study shows that effects on health caused by ASNA1 mutations can have different biochemical bases.

## INTRODUCTION

ASNA-1/GET3/ASNA1 is an essential cytosolic ATPase that is evolutionarily related to the bacterial arsenite transport factor ArsA (Kurdi-Haidar et al., 1998), however its function in eukaryotes has clearly separated (Farkas et al., 2019). The mammalian homolog ASNA1 not only plays an essential role in tail-anchored (TA) protein insertion (Colombo et al., 2016) and insulin secretion (Norlin et al., 2017) but has been characterized as an important biomarker in treatment response in schizophrenia (Mamdani et al., 2013), predicting disease severity of dengue virus infection (Soe et al., 2018), differentiation between subjects with active and latent tuberculosis (Mistry et al., 2007), a biomarker for abnormalities associated with ultra-high-risk for psychosis (Zhang et al., 2020), as well as a biomarker for Down’s syndrome (Sui et al., 2015). Mutations in ASNA1 are associated with pediatric cardiomyopathy (Verhagen et al., 2019) and mutations in its pathway protein CAML are linked in patients to hypotonia, brain abnormalities, and epilepsy (Wilson et al., 2022). Further ASNA1 interacts with and likely modulates the function of VAPB which is mutated in cases of amyotrophic lateral sclerosis (Baron et al., 2014). Therefore, it is clear that ASNA1 has a role in a variety of human diseases and a deeper understanding of clinically relevant functions of ASNA1 is of general interest.

The diverse roles of ASNA1 in different disease manifestations are reflected in the different biochemical roles of this protein. The best understood and most intensively studied the role of ASNA-1/GET3/ASNA1 is in the endoplasmic reticulum (ER) targeting of a special class of membrane proteins called tail-anchored proteins (TAP) (Chartron et al., 2012). This function strictly requires ATPase activity (Mateja et al., 2009) and the transmembrane regions of TAP proteins associate with the hydrophobic groove in dimeric ASNA1 before their insertion into the ER membrane. Insertion of TAPs via ASNA-1 requires the membrane proteins WRB/CAML (in mammals) or Get1/Get2 (in yeast) (Vilardi et al., 2014) to which ASNA1 hands off the tail-anchored proteins.

However, there are alternative roles for ASNA1/GET3 where it acts independently of its TAP insertion function and thus does not require it to interact with partners Get1/Get2 or WRB/CAML. Furthermore, Get3 is co-regulated with the proteasome/ubiquitin/NPL4 pathway and displays genetic interactions with it, while Get1 and Get2 do not. Mammalian ASNA1 interacts with the tail-anchored protein, VAPB, only via the MSP domain of the VAPB and not by its transmembrane domain. The region of mammalian ASNA1 that binds to VAPB is the FFAT motif and not its hydrophobic groove that is important in its interaction with tail-anchored proteins. Hence mammalian ASNA1 can display novel interaction patterns even with a tail-anchored protein that does not involve the canonical ASNA1/WRB/CAML pathway. Moreover, evidence suggests that VAPB may work as an alternative receptor for ASNA1 instead of WRB/CAML (Baron et al., 2014). Importantly, elegant biochemical and cell biological studies have demonstrated upon structural rearrangements GET3 acts as a holdase chaperone that binds to aggregated proteins under conditions of oxidative stress conditions and ATP depletion(Powis et al., 2013; Voth et al., 2014). This rearrangement happens after the oxidation of conserved cysteines in Get3 and converts the dimeric GET3 to a tetramer. This structural change masks the hydrophobic domain required for TA protein binding and substantially reduces its ATPase activity: Both the binding and ATPase properties of GET3, which are essential for its interaction with WRB/GET1, are completely dispensable for the function of oxidized Get3. These lines of evidence demonstrate that under conditions of oxidative stress, Get3 promotes functions that do not require WRB/GET1. Moreover, Voth et al., 2014 show that oxidized ASNA-1 also acts as a general chaperone that can refold denatured citrate synthase in vitro in the absence of WRB/GET1 (Voth et al., 2014). In *S. cerevisiae* GET3 directly binds to non-tail anchored proteins: to the chloride transporter Gef1p (Metz et al., 2006) and to the Gα subunit Gpa1 to act as a guanine nucleotide exchange factor (Lee and Dohlman, 2008). In both cases, the binding is biologically meaningful since it leads to altered biochemical outcomes that are independent of GET1.

Our work on *C. elegans* ASNA-1 showed that ASNA-1 promotes insulin secretion and this later led to the findings that insulin secretion is defective in ASNA1 knockdown mice leading to type 2 diabetes (Kao et al., 2007; Norlin et al., 2016). Further, the role of ASNA1 in cisplatin sensitivity of human tumor cells was also modeled successfully in worms, and importantly the human ASNA1 gene can replace the worm gene (Hemmingsson et al., 2010; Kao et al., 2007). Our previous work in worms revealed the TAP targeting function of ASNA-1 and showed that cisplatin exposure negatively impacts only targeting of the ASNA-1-dependent tail-anchored protein SEC-61β to the ER while having no effect on ASNA-1 independent TAPs (Raj et al., 2021).

Clinically relevant roles of ASNA-1 in insulin secretion and protection from cisplatin toxicity can be separated based on the oxidative state of the protein. As with GET3, ASNA-1 exists in two redox-sensitive states: reduced (ASNA-1^RED^) and oxidized (ASNA-1^OX^) (Raj et al., 2021). The single point mutant *asna-1(ΔHis164)*, in which the protein preferentially exists in the oxidized state, was not only as sensitive to cisplatin treatment as a complete deletion mutant but also had a severe defect in TAP insertion (Raj et al., 2021) showing that separation of function mutations exists and indicating that ASNA-1 function in insulin secretion is most likely independent of its role in TAP targeting and cisplatin response.

For a multifunctional protein with non-overlapping functions, drugs affecting one function while not perturbing the other functions would reduce the undesirable side effects and might increase chemotherapeutic response. We screened for viable and homozygous *C. elegans asna-1* point mutants to explore the possibilities of completely separating clinically relevant functions of *asna-1* with genetic variants. As a result, we characterized the highly conserved alanine in position 63 as a relevant target for function separation. *asna-1(A63V)* mutants did not display any morphology, growth, lifespan, brood size, or response to oxidative stress phenotype that is shown by *asna-1* deletion mutants. All these features made *asna-1(A63V)* animals an excellent model to study the role of *asna-1* in the response to cisplatin without the compounding effects of other phenotypic defects. Indeed, *asna-1(A63V)* mutants showed cisplatin sensitivity defect and TAP insertion defect that was as severe as that seen in deletion mutants without any detectable defect in insulin secretion or signaling pathways. Moreover, this change led to higher levels of the oxidized form of the protein and altered its subcellular distribution. We also expanded our analysis of the *asna-1(ΔHis164)* mutant to show that it has normal levels of insulin secretion and function. We performed *in silico* modeling to better understand the consequences of the structural changes caused by the single point mutations. Finally, we examined the expression of ASNA-1 and performed selective depletion to identify the tissue requirements for growth promotion. Taken together our study provides evidence that genetic or pharmacological perturbations in the Switch I and Switch II domains can selectively affect individual functions of this bifunctional protein. This study supports the notion that small molecule drugs targeting specific domains will greatly increase the cisplatin sensitivity of resistant solid tumors.

## RESULTS

### ASNA-1 is broadly expressed in *C. elegans*

Expression from an extrachromosomal transgene expressing ASNA-1:GFP (*svIs56*) was seen in sensory neurons, intestine, and hypodermis (Kao et al., 2007). Taking advantage of CRISPR/Cas9 technique we obtained a strain where two tags, mNeonGreen and AID, were inserted into the genomic locus of ASNA-1 at the C-terminus of the protein (ASNA-1::mNG::AID; *syb2249*; strain PHX2249). The mNeon Green tag allowed us to more precisely characterize the ASNA-1 expression pattern. The mNeonGreen signal was visible strongly in the pharynx, intestine, muscles, proximal and distal germline cells, oocytes, developing vulva, vulval muscles, spermatheca, sperm, sheath cells, the most proximal pair of gonadal sheath cells (**Fig. S1**), and in head neurons (**Fig. S2**). This analysis not only confirmed our previous observations (Kao et al., 2007) but also characterized previously unknown ASNA-1 expression sites in the *C. elegans* reproductive system. We concluded that germline and somatic gonad are likely important places for ASNA-1 function given the broad expression of ASNA-1 in this tissue. To more precisely identify the cells expressing ASNA-1 in each tissue we next examined a bi-cistronic ASNA-1^SL2::mNeonGreen::H2B allele *syb5730* of the gene (strain PHX5730). In this construct, the ASNA-1 mRNA independently makes untagged ASNA-1 and nuclear-localized mNeonGreen to report on each nucleus that expresses the *asna-*1 mRNA. Analysis of the mNeonGreen signal in *asna-1(syb5730)* animals showed that virtually every cell in the body expresses the ASNA-1 mRNA throughout development in both the soma and the germline (**Fig. S14**).

### Soma-specific *asna-1* knockdown leads to L1 larval arrest

Worms lacking maternal and zygotic *asna-1* arrest at the 1^st^ larval (L1) stage even in the presence of food (Kao et al., 2007). L1 arrest is a characteristic of wild-type larvae hatched in the absence of food or of strong *daf-2*/insulin receptor mutants (Gems et al., 1998). Taking into consideration how broadly ASNA-1 is expressed in somatic and germline tissues, we asked whether somatic depletion of ASNA-1::mNG::AID would cause a larval arrest phenotype similar to that seen after complete *asna-1* genetic depletion. Taking the advantage of the auxin-inducible tissue-specific knockdown system, we created the strain expressing ASNA-1::mNG::AID (*syb2249*) and TIR1 pan-somatic driver *ieSi57. Arabidopsis* TIR1 is an F-box protein that interacts with the AID tag in the presence of auxin to assemble an E3 ubiquitin ligase complex and targets AID-containing proteins for rapid degradation by the proteasome (Ashley et al., 2021; Zhang et al., 2015). Tissue or cell-type-specific expression of TIR1 limits the degradation to of AID tagged proteins to TIR1 expressing cells. This system allowed us to assess the depletion of ASNA-1 in all somatic tissues using the pan-soma TIR1 driver *ieSi57*. 4th larval stage (L4) hermaphrodites were exposed to 1mM auxin (AUX) for 48h while they produced progeny and then removed from the plate. Their progeny remaining on the auxin-containing plates were analyzed 48 hours after removal of the mothers (**Fig. 1**). Similarly handled L4 worms on non-auxin NGM plates served as a control. First, we observed that *asna-1::mNG::AID*;*ieSi57* mothers did not display any morphological defects after 48h exposure to auxin but did produce fewer progeny in comparison to non-treated control animals. On auxin plates all their progeny arrested at the L1 stage (**Fig. 1**). The auxin-mediated depletion of ASNA-1 was effective because the mNeon green signal was not visible in any of the previously characterized somatic tissues (**Fig. 1**). However, we saw a strong mNeonGreen signal in the Z2 and Z3 germline progenitor blast cells (**Fig. 1**), which give rise to the entire germline during development (Kimble and Hirsh, 1979). None of the arrested larvae, even 72 hours after removal of the mothers, showed more than the two cells (Z2 and Z3) in the gonad primordium. The cells could be identified as Z2 and Z3 because they lay in the center of the gonad primordium, were in contact with each other (Kimble and Hirsh, 1979), and expressed ASNA-1::mNG::AID while it was depleted in all surrounding somatic cells. To further characterize the effects of ASNA-1 depletion, we sought to establish if *asna-1::mNG::AID*;*ieSi57* worms exposed to auxin only from the 1st larval stage onwards would display further development and possibly reach later larval stages. Staged L1 hermaphrodites were exposed to 1mM auxin (AUX) for 24h, 48h, and 72h and analyzed at these time points. (**Fig. S3**). Similarly staged unexposed *asna-1::mNG::AID*;*ieSi57* larvae served as controls. We observed that all animals on AUX plates were still arrested at the L1 stage (**Fig. S3, Fig S4**), whereas control animals reached adulthood. As expected, auxin-treated animals were characterized by a lack of ASNA-1::mNG::AID expression in the somatic tissues and we observed a strong ASNA-1::mNG::AID signal in the germline progenitors (**Fig. S4**). However, in contrast to worms hatched on AUX-containing plates, ASNA-1::mNG::AID was seen in many more germline cells with a maximum of 14 positive cells (**Fig. S4**). In total, 13/20 worms analyzed had more than 4 germline cells. This was in contrast to the mNeonGreen signal only being observed in the two primordial germ cells Z2 and Z3 in animals hatched on AUX plates (**Fig. 1**). However, the larvae in the second experiment did not have the alae that are characteristic of L2 animals indicating that depleting somatic ASNA-1::mNG::AID after the birth of L1 larvae allowed for further development of the germline but did not allow the larvae to reach the L2 larval stage. The size of the gonad was also characteristic of L1 stage larvae. By contrast, auxin-mediated depletion of ASNA-1 from the L4 stage onwards did not block progression to the adult stage.

**Figure 1.**
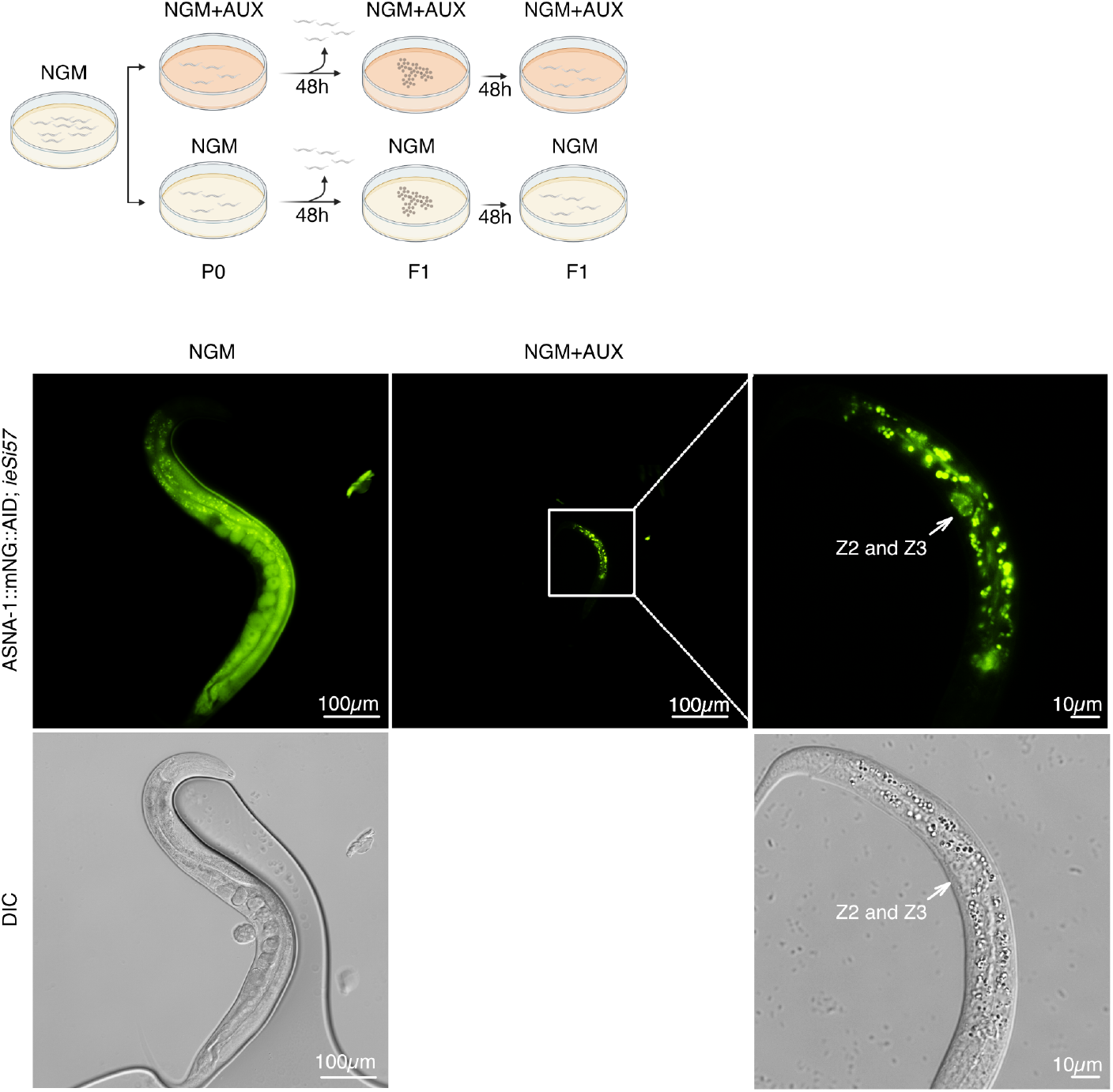
Soma-specific ASNA-1 depletion leads to L1 larval arrest. Schematic representation of first experimental setup for auxin-inducible pan-somatic ASNA-1 depletion. Representative fluorescent and differential interference contrast (DIC) images of worms expressing ASNA-1::mNeonGreen::AID (*syb2249*) and pan-somatic TIR1 driver *ieSi57* grown on plates without (NGM) or with (NGM+AUX) 1mM auxin. White arrows indicate the Z2 and Z3 cell positions.

### Intestine-specific *asna-1* knockdown leads to developmental delay

Our previous research has established that *asna-1* is required cell non-autonomously for growth. Both neuron and intestine-specific promotors driving *asna-1* expression were able to rescue the *asna-1(ok938)* body size phenotype (Kao et al., 2007), while ASNA-1 is required in the intestine but not the neurons for the rescue of the cisplatin sensitivity phenotype (Hemmingsson et al., 2010). We have also seen a requirement for ASNA-1 in the gut for proper tail-anchored protein targeting (Raj et al., 2021). Therefore, we sought to establish a role of the intestinal function of ASNA-1 in growth and development to ask whether the L1 arrest phenotype could be ascribed to the TAP targeting function in the intestine. For that purpose, we constructed a strain expressing an intestine-specific TIR1 driver *ieSi61* in the ASNA-1::mNG::AID (*syb2249*) background. To compare pan-somatic depletion of ASNA-1 with gut-specific depletion, as before we exposed *asna-1::mNG::AID;ieSi61* L4 hermaphrodites to 1mM auxin (AUX) for 48h, removed them from the plate, and analyzed their progeny 48h after the removal of the mothers (**Fig. 2**). Worms on NGM plates not containing auxin served as a control. Microscopy revealed an absence of mNeonGreen fluorescence in the intestine while pharyngeal and germline expression was visible at normal levels (**Fig. 2**). Intestine-specific ASNA-1 depletion starting in developing embryos led to a significant delay in development in comparison to control animals (**Fig. 2**). The larvae hatched on these plates took an additional day to reach adulthood. When L1 larvae were exposed to 1mM AUX, no delay in growth was seen (**Fig. S5**). Thus, there was a requirement for ASNA-1 in the intestine for growth but the requirement was not as stringent as that seen with pan-somatic depletion. Based on that we concluded that growth and development dependent on ASNA-1 was a multi-tissue requirement or that auxin-mediated depletion was not complete.

**Figure 2.**
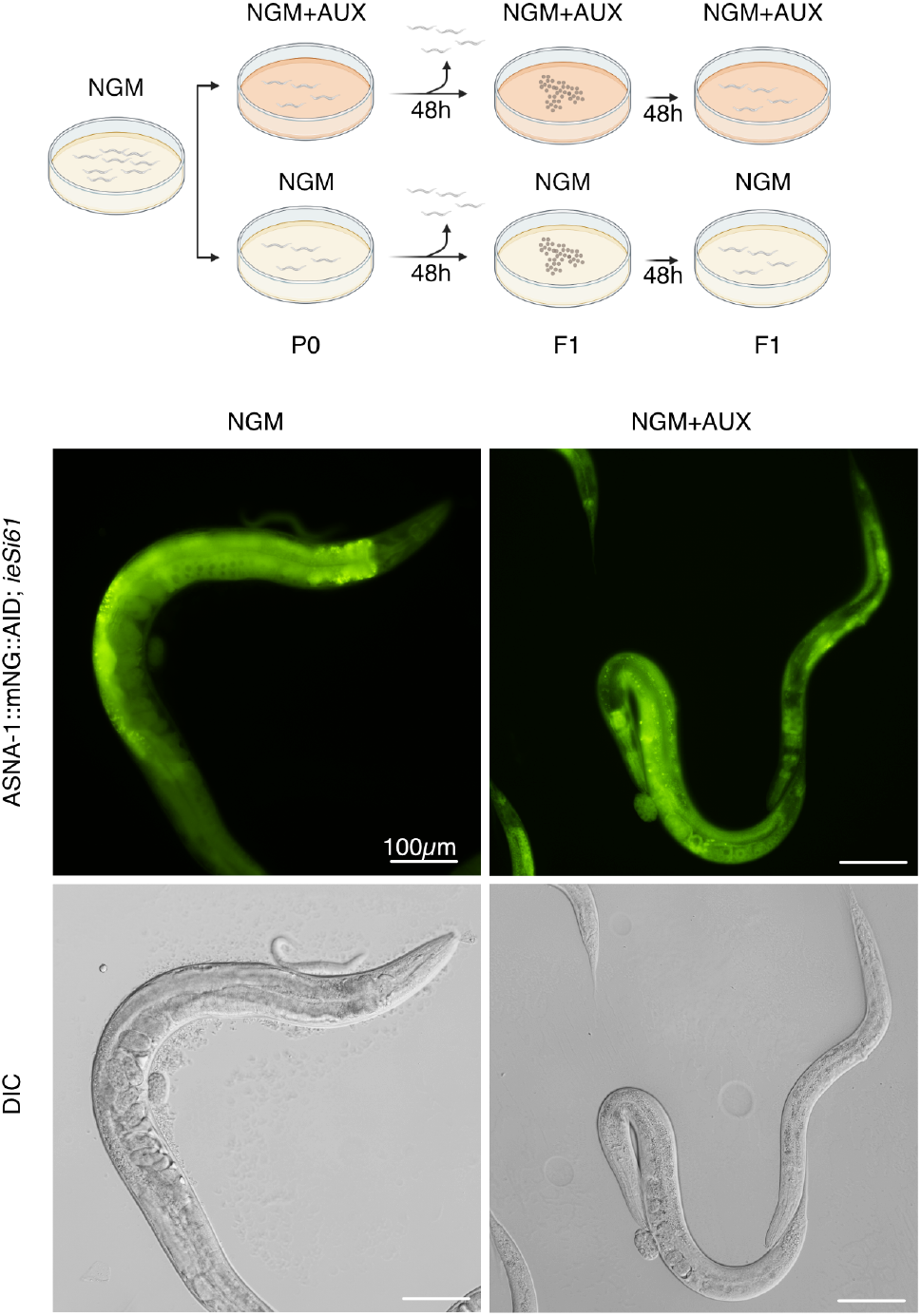
Intestine-specific ASNA-1 depletion leads to a developmental delay. Schematic representation of first experimental setup for auxin-inducible gut ASNA-1 depletion. Representative fluorescent and differential interference contrast (DIC) images of worms expressing ASNA-1::mNeonGreen::AID (*syb2249*) and TIR1 pan-somatic driver *ieSi61* grown on plates without (NGM) or with (NGM+AUX) 1mM auxin.

### Simultaneous depletion of ASNA-1 in the intestine and neurons is equivalent to pan-somatic depletion

Having shown that intestinal depletion of ASNA-1 was not sufficient to produce the L1 growth arrest phenotype, we wished to determine whether we could narrow down the tissue requirements that lead to a strong L1 arrest phenotype seen in *asna-1* mutants (Kao et al., 2007) upon pan-somatic depletion. To do this we first depleted by auxin treatment ASNA-1::mNG::AID in neurons using the pan-neuronal driver *reSi7* (Ashley et al., 2021). *asna-1::mNG::AID (syb2249)*; *reSi7* worms were exposed to auxin from the time of hatching by putting 4th larval stage mothers on auxin-supplemented NGM plates. The worms produced a normal number of progeny and the progeny displayed no growth defects or developmental delays. To ask whether the 1st larval stage arrest was caused by simultaneously depleting ASNA-1 in both the intestine and neurons, we next made and tested a strain of the genotype *asna-1::mNG::AID (syb2249);reSi7;ieSi61*. In this strain, the two TIR1 drivers drove the depletion of ASNA-1 in both tissues. In these worms, we observed that when 4th larval stage mothers were allowed to produce progeny on auxin-supplemented NGM plates, they still produced a large number of progeny but all the progeny (n>250) arrested as 1st larval stage sized animals. This reproduced the pan-somatic ASNA-1 depletion phenotype and indicated that both a neuropeptide secretion defect (such as DAF-28 secretion) and a defect in the trafficking of SEC-61β alone or along with other TAPs is required for the control of growth at the 1st larval stage (**Fig. S15**).

### Germline-specific *asna-1* knockdown does not play a role in driving ASNA-1 functions

Extensive research has shown the importance of communication between soma and germline (Gaddy et al., 2021; Kishimoto et al., 2017; Nono et al., 2020; Qi et al., 2021). Based on our somatic and intestinal ASNA-1 depletion experiments, our observation of extensive ASNA-1 expression in the germline as well as the sterility phenotype of *asna-1* deletion mutants, we asked if germline-specific knockdown of ASNA-1 in larval or adult stage would lead to the same phenotype as seen in strain with somatic tissue knockdown. In the same fashion, we created the strain expressing ASNA-1::mNG::AID (*syb2249*) and TIR1 germline-specific driver *ieSi64*. This allowed us to assess the depletion of ASNA-1 only in the germline, leaving the somatic expression of ASNA-1 unchanged. As before, L4 larvae were exposed to 1mM auxin (AUX) for 48h, removed from the plate and their progeny were analyzed after 48h after the removal of the mothers (**Fig. S6A**). Worms on non-auxin NGM plates served as a control. First, mothers exposed to AUX laid fewer progeny in comparison to non-auxin controls (**Fig. S6A**). Second, the progeny that hatched reached adulthood 48h after removal of mothers, at the same rate as the non-auxin treated control worms (**Fig. S6A**). Moreover, there was no visible developmental or growth defect when L1 larvae were exposed to auxin (**Fig. S6B**). We concluded that depletion of ASNA-1::mNG::AID in germline using this TIR1 driver largely did not affect the development of the worms and did not cause larval arrest as in the case of the pan-somatic depletion of ASNA-1, but affected reproduction and progeny formation. However, using the transgenic system available the effect of ASNA-1 germline depletion on reproduction was not as severe as that seen in genetic deletion mutants. Taken together, this indicated that ASNA-1 expression in the somatic tissue of *C. elegans* played a major role in driving ASNA-1 functions.

### Insulin signaling is normal and germline development defects are less severe in *asna-1(ΔHis164)* compared to *asna-1* deletion mutants

Our previous research showed that *asna-1*(*ΔHis164)* mutants were as sensitive to cisplatin as deletion mutants in the gene and have a severe TAP insertion defect (Raj et al., 2021). Further, we showed that the ASNA-1^ΔHis164^::GFP protein largely exists in the oxidized state (Raj et al., 2021). Transgenes expressing *asna-1*^*ΔHis164*^::GFP rescued the *asna-1* growth phenotype of the deletion mutant indicating indirectly that the point mutant protein can restore insulin signaling (Hemmingsson et al., 2010). To examine the growth phenotypes of *asna-1(ΔHis164)* mutants, we directedly measured the insulin secretion and signaling efficiency by analysis of DAF-16::GFP localization and DAF-28/insulin::GFP secretion. The cytoplasmic localization of DAF-16::GFP indicates high IIS levels since DAF-16::GFP is localized to nuclei in mutants with insulin signaling defects (Henderson and Johnson, 2001). DAF-28/insulin::GFP secreted from neurons and taken up by the scavenger cells called coelomocytes reports on insulin secretion (Kao et al., 2007). In *asna-1(ΔHis164)* animals DAF-16::GFP protein was localized to the cytoplasm whereas the DAF-28/insulin::GFP protein was secreted and taken up by coelomocytes at normal levels (**Fig. 3A**). Both markers directly show that insulin secretion and signaling were unaffected in *asna-1(ΔHis164)* mutants in comparison to the strong defects in *asna-1* deletion mutants. Although both mutants did differ in IIS status, *asna-1(ΔHis164)* animals displayed a sterility phenotype as did *asna-1* deletion mutants. The bigger size of the *asna-1(ΔHis164)* worms (**Fig. 3B**) suggested that germline defects in both mutants differ significantly. We sought to understand in detail these differences by examining the germline phenotype of *asna-1(ok938)* and *asna-1(ΔHis164)* animals. For this purpose, we used the transgenes to simultaneously monitor germ cell membranes and germline nuclei *in vivo. asna-1(ok938)* animals do not produce any oocytes or sperm and show a gonad migration defect (**Fig. S7A**). *asna-1(ΔHis164)* also showed a gonad migration problem but the germlines contained oocytes and sperm as well as non-fertilized embryo-type structures in the uterus. These cells contained only one nucleus and displayed no embryonic divisions (**Fig. S7A, B**). Ovulation, the transit of the ovum through the spermatheca, and entry of oocytes into the uterus require the presence of sperm (McCarter et al., 1999). This led us to investigate whether *asna-1(ΔHis164)* mutants might be defective in fertilization, taking into consideration the strong ASNA-1 expression in sperm and spermatheca in wild-type worms (**Fig. S1, Fig. S14**). Indeed, the *itIs37* marker expressing *pie-1p::mCherry::his-58* to visualize germline nuclei showed that sperm was present in the spermatheca in *asna-1(ΔHis164)* animals as in *asna-1(+)* (**Fig. S7B**). The fact that some oocytes could complete ovulation and reach the uterus implied that immature spermatids or primary spermatocytes were present in *asna-1(ΔHis164)* mutants. However, since the “post-oocyte”, cells did not contain more than one nucleus, we concluded that there was a defect either in oocytes or sperm of *asna-1(ΔHis164)* animals. The difference in germline defect between point mutants and deletion mutants stands in striking contrast to the similarity in cisplatin sensitivity phenotype of these two mutants (Raj et al., 2021) indicating that different functions can be independently targeted.

**Figure 3.**
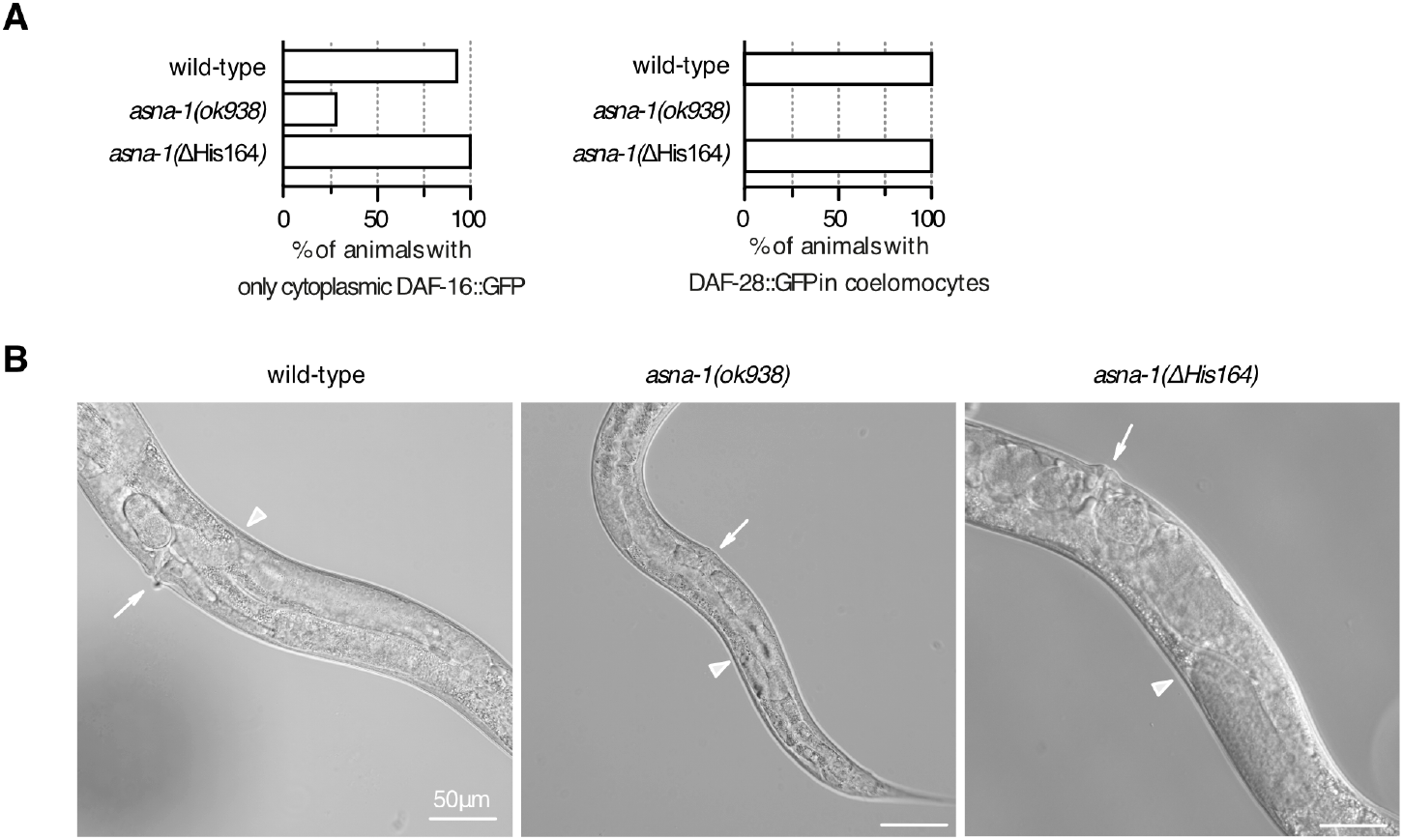
Comparison of germline and insulin/IIS defects *in asna-1(Δ His164)* and *asna-1*(*ok938)* deletion mutants. (A) Percentage of 1-day old adults with indicated genotypes displaying only cytoplasmic localized DAF-16::GFP (n ≥ 15) and percentage of 1-day old adults with secreted DAF-28::GFP in coelomocytes (n ≥ 20). (B) Representative DIC pictures of adults showing the body size of wild type, *asna-1(ok938)*, and *asna-1(ΔHis164)* strains.

### Analysis of ASNA-1 point mutants reveals the importance of a conserved alanine for cisplatin resistance

We have previously shown that *C. elegans* ASNA-1 exists in two redox states: reduced (ASNA-1^RED^) and oxidized (ASNA-1^OX^), and that important biological functions of the protein are separable based on the redox state (Raj et al., 2021). As mentioned, these conclusions were drawn based on an analysis of the *asna-1* single point mutant: *asna-1(ΔHis164)*, which also showed the sterility phenotype described above. Therefore, we wondered if this was a unique characteristic of mutating His164 or whether any other single point mutation in the ASNA-1 protein would also be able to completely separate the important biological functions of ASNA-1: insulin secretion, TAP targeting, and cisplatin sensitivity while not causing sterility in the worm. For this reason, we examined a set of seven strains generated by the Million Mutation Project, each carrying a missense point mutant in ASNA-1 along with other mutations in the genome (Thompson et al., 2013) (**Table S1**). Of these seven, only in four mutants were the mutated amino acids conserved with human ASNA1 (**Table S1; Fig. S6A**). None of the single amino acid changes caused reduced steady-state levels of the ASNA-1 protein (**Fig. S9**). We performed a cisplatin survival analysis of all seven strains to ask if any of those mutants could increase sensitivity to cisplatin treatment. The LD_50_ of cisplatin for *asna-1(ok938)* deletion mutants was 300 µg/mL after 24h treatment (Hemmingsson et al., 2010), and hence we used this concentration for our experiments. We found that only two strains, *asna-1(gk687101)* and *asna-1(gk592672)*, showed increased cisplatin sensitivity after 24h exposure to the drug (**Fig. 4A**). Since *asna-1(gk592672)* strain produced a more severe sensitivity phenotype, we focused on this point mutation in our further analysis. The *asna-1(gk592672)* mutation is an alanine to valine change, at a highly conserved position 63 which corresponded to alanine 82 in the human homolog. This amino acid is positioned close to the Switch I domain. This domain plays an essential role in the activation of ATP hydrolysis during TA protein targeting (**Fig. S8A**). We used the GnomAD browser (https://gnomad.broadinstitute.org/) to assess the effect of this point mutation in the human homolog of ASNA-1. The browser provides not only a list of variations but also a critical view of the clinical application (Lek et al., 2016). In the exome sequence data for 60,706 individuals, ASNA1 has 79 observed single nucleotide variants (SNVs) that resulted in missense mutations. In those, A82 was detected to be mutated to threonine (T) (A82T) with two allele counts in the European (non-Finnish) population. PoyPhen-2 (http://genetics.bwh.harvard.edu/pph2/) is a tool that predicts the possible impact of amino acid substitution on the structure and function of human protein. The human ASNA1(A82T) mutation was classified as ‘probably damaging’. This was also true for the alanine to valine substitution. Mindful of other mutations in other genes in the *asna-1(gk592672)* containing strain we outcrossed the strain 13 times and performed the cisplatin sensitivity assay again. Indeed, the outcrossed strain, hereafter called *asna-1(A63V)*, was as sensitive to cisplatin exposure as the *asna-1*(*ok938)* deletion mutant. The cisplatin sensitivity phenotype was rescued by a single copy of wild-type ASNA-1 fused to GFP (*knuSi184*) expressed from a transgene (**Fig. 4B**). This transgene also rescued the ASNA-1 protein null mutant for the cisplatin sensitivity phenotype (**Fig. 4B**). We concluded that change in a highly conserved alanine at position 63 was the genetic locus causing cisplatin sensitivity as severe as the deletion mutant and no other mutation in the genetic background contributed to the phenotype.

**Figure 4.**
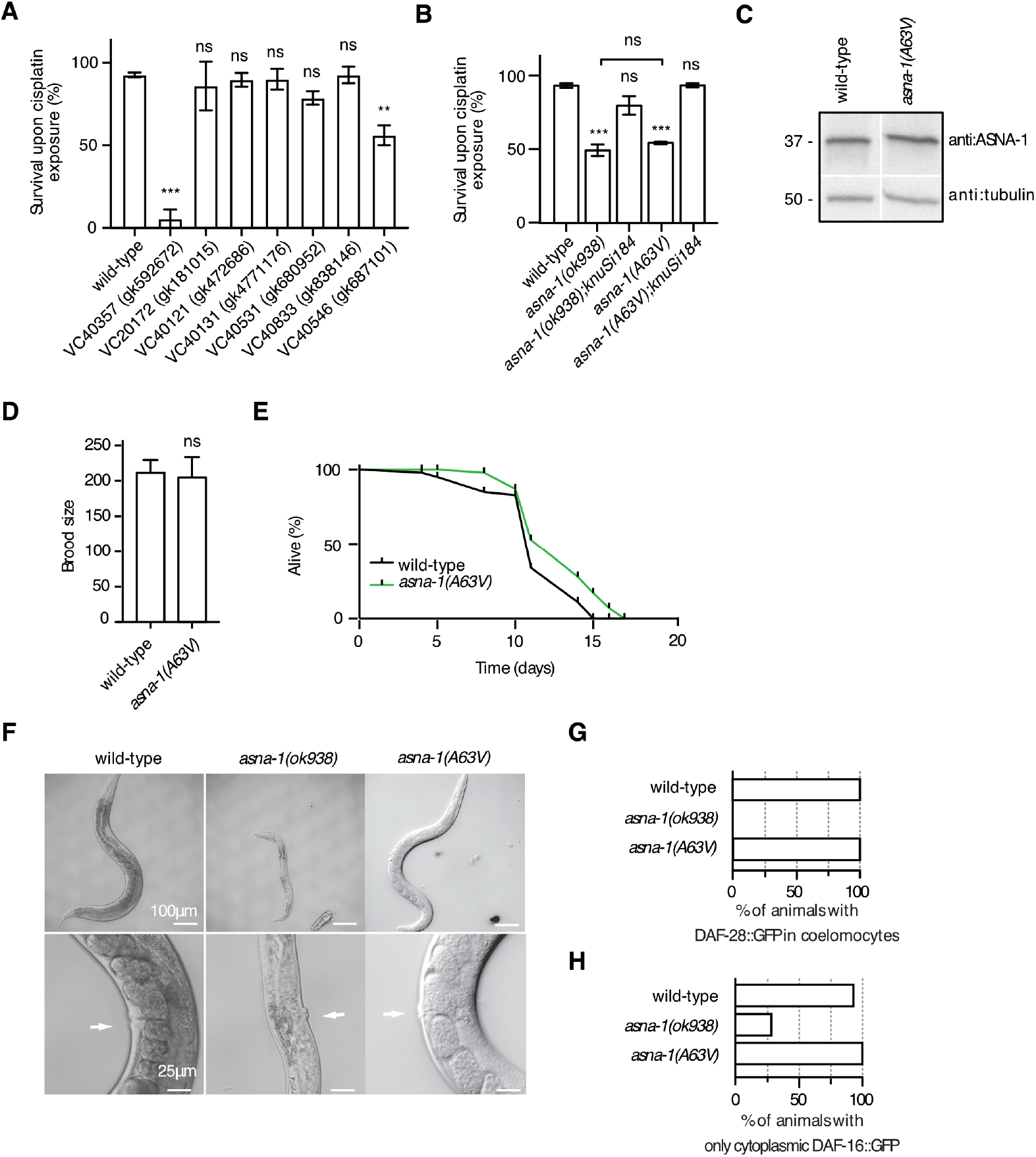
Analysis of ASNA-1 point mutants reveals the importance of a conserved alanine for cisplatin resistance. (A) Analysis of cisplatin sensitivity phenotype of seven million mutation project strains bearing point mutations in *asna-1*, (B) Rescue of the cisplatin sensitivity phenotype of *asna-1(ok938* deletion) mutants and *asna-1(gk592672* /A63V) mutants with a single copy ASNA-1(+) transgene *knuSi184*. Bars represent mean survival ± SD of 1-day-old adult animals exposed to 300 µg/mL of cisplatin for 24h. Statistical significance was determined by the one-way ANOVA followed by Bonferroni post-hoc correction (n≥50). Survival experiments were performed in triplicate. (C) Western blot analysis to estimate ASNA-1 levels in wild-type and *asna-1(A63V)* mutants. The blot was probed with an anti-ASNA-1 antibody. Tubulin was used as a loading control. For full uncropped western blot image, see Fig. S12. (D) Brood size analysis of wild-type and *asna-1(A63V)* mutants (n=5). Statistical significance was determined by the independent two-sample t-test. Bars represent mean ± SD. (E) Life span analysis of wild-type (n=40) and *asna-1(A63V)* (n=40) animals. (F) Top panel shows representative DIC pictures of adults to show the body size of wild type, *asna-1(ok938)*, and *asna-1(A63V)* strains. The magnification of the germline of the animals is shown in the bottom panel. White arrows indicate the vulva position for orientation purposes. (G) Percentage of 1-day old adults of the indicated genotypes with secreted DAF-28::GFP accumulating in coelomocytes (n ≥ 20). (H) percentage of 1-day old adults of the indicated genotypes displaying only cytoplasmic localized DAF-16::GFP (n ≥ 20).

### *asna-1(A63V)* mutants have a normal brood size and insulin/IIS signaling and secretion

To obtain more evidence for separable functions of ASNA-1, we characterized the *asna-1(A63V)* strain. As in the case of the *asna-1(ΔHis164)* protein, the A63V mutation did not affect the stability of the protein (**Fig. 4C; Fig. S12**). ASNA-1^A63V^::GFP expressed from a multi-copy transgene was detected in the ER (**Fig. S8B**) in the same manner as the wild-type protein ASNA-1::GFP. *asna-1(A63V)* animals did not display the enhanced ER stress phenotypes shown by the *asna-1(ok938)* deletion mutant (**Fig. S8C**). While *asna-1(A63V)* animals were as sensitive to cisplatin as the deletion mutant (**Fig. 4B**), they were fertile with a normal brood size and life span (**Fig. 4D, E**), and had properly developed germlines (**Fig. 4F**). This characteristic opened up the possibility of separating the cisplatin, TAP insertion, and insulin functions of ASNA-1 without compromising on the sterility phenotypes seen in *asna-1(ok938)* or *asna-1(ΔHis164)* mutants (Raj et al., 2021). Importantly, *asna-1(A63V)* mutants had neither the insulin signaling nor the insulin secretion defect (**Fig. 4G, H**), which was measured respectively by cytosolic/nuclear presence of DAF-16::GFP and uptake of DAF-28::GFP by the coelomocytes. This evidence led to the conclusion that indeed functions of ASNA-1 in cisplatin sensitivity and insulin signaling/secretion were completely separable in animals without the compromised function in the germline and further that the cisplatin sensitivity of *asna-1* mutants was purely a soma-specific defect since the germline is normal in *asna-1(A63V)* mutants.

### *asna-1(A63V)* mutants separate the TAP insertion function from insulin secretion

We have previously shown that *asna-1(ok938)* and *asna-1(ΔHis164)* mutants displayed a strong TAP targeting defect (Raj et al., 2021). This conclusion was supported by three independent analyses. Firstly, Pearson colocalization analysis following confocal microscopy of worms co-expressing the TAP, GFP::SEC-61β with the ER marker, mCherry::SP12. Secondly, N-linked glycosylation analysis of SEC-61β to monitor insertion of SEC-61β into the ER membrane. Thirdly, an analysis of steady-state protein levels of SEC-61β in *asna-1* deletion mutants, which suggested that failure of insertion into the ER results in protein degradation (Raj et al., 2021). In *asna-1(ok938)* mutants, the amount of N-linked glycosylated SEC-61β protein remained unchanged whereas the total protein level was greatly decreased (Raj et al., 2021). Since *asna-1(ok938)* and *asna-1(ΔHis164)* mutants share the cisplatin sensitivity phenotype of *asna-1(A63V)* mutants, we therefore wondered if *asna-1(A63V)* animals could support ASNA-1-dependent TAP insertion. The Pearson correlation analysis of the two co-expressed transgenes (GFP::SEC-61β and mCherry::SP12) in *asna-1(A63V)* mutants revealed a significantly defective TAP targeting phenotype (**Fig. 5A, B**). We next measured the direct insertion of the SEC-61β protein into the ER membrane (Raj et al., 2021). We used the transgene (*rawEx64)* expressing SEC-61β in which the beta-opsin tag is inserted at the extreme C-terminus and contains a reactive arginine glycosylated by ER luminal enzymes when the C-terminus is exposed to the ER lumen. This reports on the proper insertion of the SEC-61β protein into the ER membrane. The glycosylation event is revealed by the change in mobility of the glycosylated tagged protein. This analysis revealed that targeting of the TAP SEC-61β was decreased in the *asna-1(A63V)* mutant background (**Fig. S10A**) while the steady-state level of the SEC-61 β protein remained unchanged (**Fig. S10B**).

**Figure 5.**
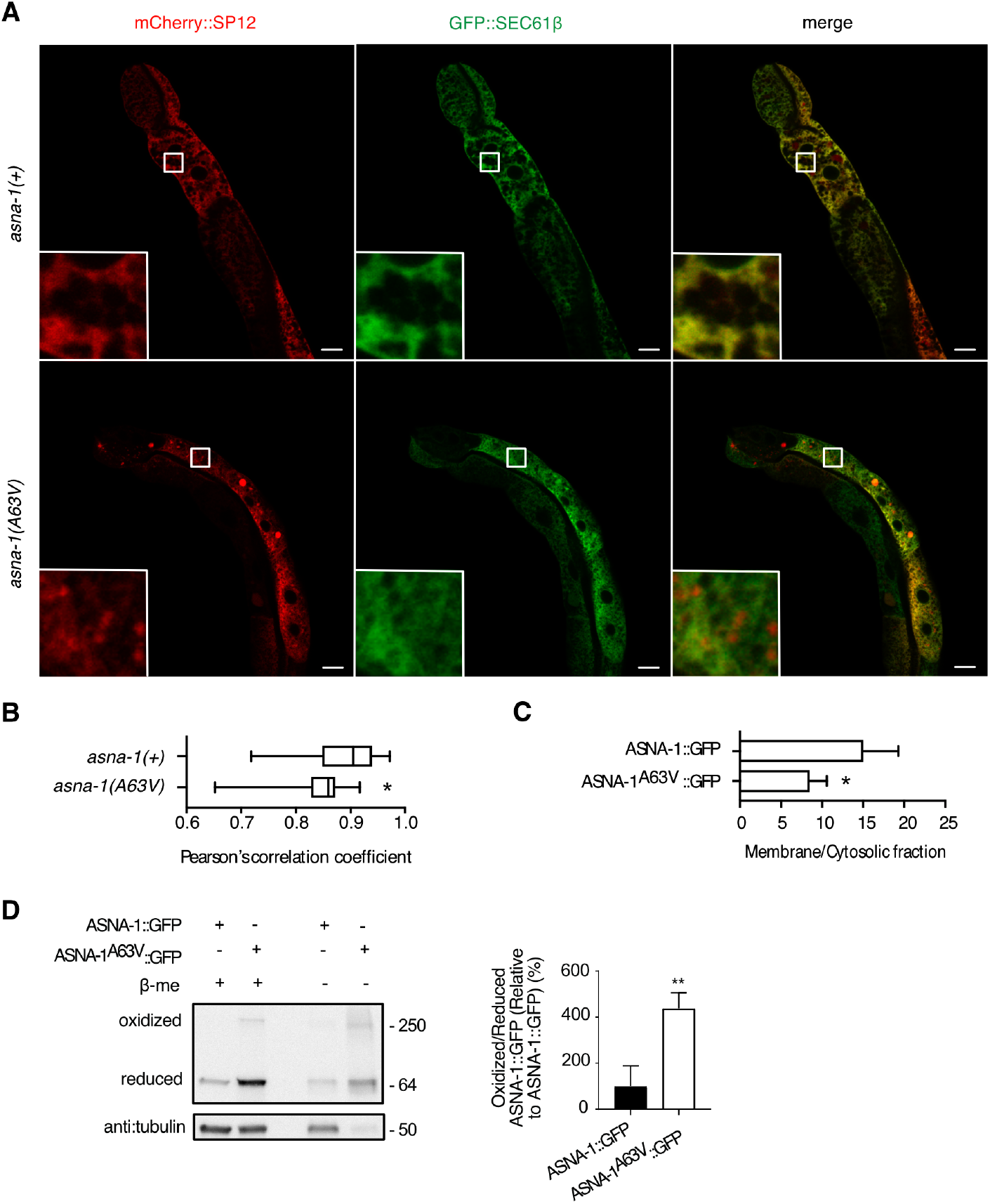
*asna-1(A63V)* mutants are defective for TAP insertion but have normal insulin secretion. (A) Representative confocal images of *asna-1(+)*, and *asna-1(A63V)* 1-day old adult animals co-expressing GFP::SEC-61β and mCherry::SP12 in the intestine. Imaging was performed in int8 and int9 cells. Scale: 10 μm and 60 μm for magnification. (B) Pearson’s correlation analysis of GFP::SEC-61β and mCherry::SP12 co-localization in different strains. The box plot represents the average Pearson correlation coefficient (R) of the indicated strains. Statistical significance was determined by the Mann-Whitney test (n ≥ 10). (C) Band intensity quantification of membrane/cytosolic fraction of ASNA-1::GFP and ASNA-1^A63V^::GFP based on the western blots presented in Figure S13A. Statistical significance was determined by the independent two-sample t-test. Error bars represent ± SD. (D) Representative western blot after reducing and non-reducing SDS-PAGE to detect levels of oxidized and reduced ASNA-1^A63V^::GFP. Control worms expressed the ASNA-1(+)::GFP transgene. Blots were probed with anti:GFP antibody and tubulin was the loading control. For full uncropped blot source images including replicates used for quantification, see Fig. S13B. Band intensity quantification of oxidized/reduced ASNA-1::GFP. Statistical significance was determined by the independent two-sample t-test. Experiments were performed in triplicate. Bars represent ± SD.

### The A63V mutation alters the subcellular localization of ASNA-1

ASNA-1 physically interacts with the ER membrane resident receptor WRB-1 (Raj et al., 2021). We next investigated whether the defect in GFP::SEC-61β targeting might be also associated with the reduced association of the ASNA-1(A63V) protein with the membrane. Separation of cytoplasmic from membrane fractions by ultracentrifugation revealed significantly reduced membrane association of the ASNA-1^A63V^::GFP protein compared to that seen in for ASNA-1^+^::GFP (**Fig.5C, Fig. S13**). We concluded that *asna-1(A63V)* mutants shared the TAP insertion defect seen in the *asna-1* deletion mutants likely via a reduced ability to bind to ER membranes. This defect was manifested without affecting the insulin signaling and indeed those two vital functions of ASNA-1 were likely independent of each other.

### The ASNA-1(A63V) mutant protein preferentially exists in the oxidized state

We have previously shown that *C. elegans* ASNA-1 is a redox-modulated protein and its functions are separable based on the redox state of the protein (Raj et al., 2021). We have obtained evidence for this functional separation based on the analysis of *asna-1(ΔHis164)* point mutants in which the protein preferentially exists in an oxidized state. This mutant was defective for TAP insertion and cisplatin resistance (Raj et al., 2021) and as shown in Figure 3, these mutants were normal for insulin secretion/signaling function. However, as mentioned above, *asna-1(ΔHis164)* although bigger in size than deletion mutants (**Fig. 3B)**, showed a sterility phenotype. We therefore wondered, based on the analysis of *asna-1(A63V)* mutants, if *asna-1(A63V)* animals the separation of vital ASNA-1 functions are reflected in changes in the oxidative state of the protein. To address this, we performed non-reducing SDS-PAGE to evaluate the oxidative state of *asna-1(A63V)* animals. Indeed, the analysis revealed that four fold more of the ASNA-1(A63V) protein exists in the oxidized state (**Fig. 5D, Fig. S13**). Strikingly, the shift in ASNA-1^A63V^::GFP to the oxidized state occurred in the absence of an increased expression of markers of ER stress (*hsp-4*) (**Fig. S8C, D**), or mitochondrial stress markers (*hsp-6* and *hsp-60*) (**Fig. S8E**), whereas oxidative stress markers (*gst-4, gst-30*, and *gst-38*) were slightly increased (**Fig. S8F**). We concluded that the A63V mutants in ASNA-1, as previously seen in ΔHis164 point mutants, had inherently high ASNA-1^OX^ levels, which resulted in sensitivity to cisplatin. They also had a TAP insertion defect but maintained not only normal insulin secretion function but also did not have a compromised germline phenotype.

### *in silico* modelling of ASNA-1(A63V) and ASNA-1(ΔHis164) proteins predicts the importance of Switch I and Switch II domains in function separation

The model of the wild-type ASNA-1 protein assembled as a dimer shows that the dimerization interface presents three main interaction regions: the Zinc atom coordination, the ATP binding site, and the Histidine164 contacts (**Fig. S11**). The model obtained by the single point mutation A63V showed that the substitution of alanine by valine reduced the distance between the helix α3 and the helix α2 by the addition of the isopropyl group at the position 63 (helix α3) pointing directly towards the Thr32 (helix α2) in the P-loop motif (**Fig. 6A**). The model proposed a movement correction of the threonine hydroxyl group induced by this closed contact with the Valine. The Thr32 plays a key role in the coordination of the Mg atom (**Fig. 6A, Fig. S11**), its hydration state, and thereby the binding and catalysis of the ATP. The binding of ATP is a key event in the formation of the closed dimer and in the formation of the closed interface by its direct participation in it. We conclude that A63V change might influence the P-loop structure and affect ATP binding.

**Figure 6.**
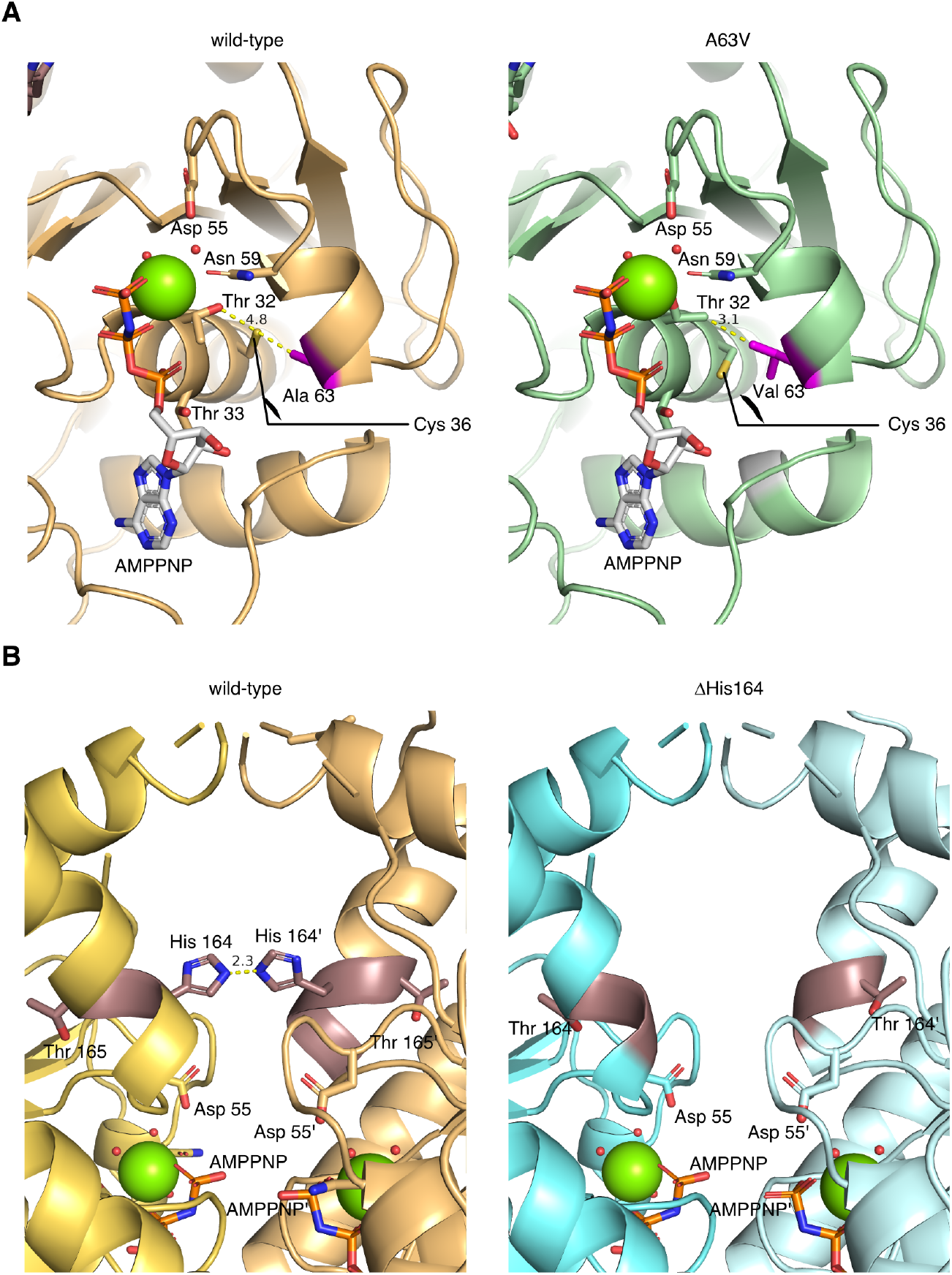
Structural representation of the effect of the A63V and ΔH164 mutations in the ATP-bound ASNA-1 dimer complex. The *in-silico* models of *C. elegans* ASNA-1 mutants show the structural impact of these mutations in the key processes of ASNA-1 ATPase activity and the dimer complex formation. (A) In the left panel, the model of an ASNA-1 wild-type protein monomer in complex with Mg (green) and AMPPNP (white) is shown in light orange. The amino acid Ala63 is shown in pink. The amino acids depicted show the influence of the amino acid at position 63 on the amino acids coordinating the Mg atom. In the right panel, the monomer of ASNA-1(A63V) is represented in green and the Val63 in pink. The distances between the amino acid at position 63 and Thr32 are shown in yellow. (B) In the left panel is represented each monomer of the ASNA-1 wild-type dimer in complex with Mg (green) and AMPPNP in light orange and yellow-orange, in deep salmon is shown the position of His164. The distance between the His164 of each monomer is displayed in yellow. In the right panel, each monomer of ASNA-1(ΔHis164) is colored in cyan and light cyan, and the amino acid Thr164 is colored in dark salmon.

The wild-type model showed the close interaction between the His164 and His164’ of each monomer to perform the dimer interface (**Fig. 6B**). The model of ASNA-1 presenting the deletion of the His164 showed that the loss of histidine implies a loss of one of the key amino acids that create the dimeric interface and does not suggest the formation of compensatory interactions. (**Fig. 6B**). Therefore, the deletion of this amino acid would affect the stability of the closed conformation and probably its formation in a negative fashion.

## DISCUSSION

An intensively studied role of ASNA-1 homologs is in the targeting of a subset of tail-anchored proteins to the ER membrane and beyond. However, it has also been clear that ASNA-1 and its homologs have roles that are not connected to its tail-anchored targeting function. These include chloride channel function, a holdase function for aggregated proteins, and the binding to the ALS-related protein VAPB protein among others (Baron et al., 2014; Powis et al., 2013; Schmidt et al., 2005). Earlier work from our lab has also addressed this issue and separated insulin secretion from cisplatin response (Raj et al., 2021). A specific objective of this study was to examine further the relative contributions of different tissue in worms to functions related to growth, reproduction, and cisplatin sensitivity. A second major goal was to discover whether single point mutations in *C. elegans* ASNA-1 will fulfill the requirements for ASNA-1 function of cisplatin response separate from its roles in growth and reproduction.

First, we characterized the expression pattern of ASNA-1 in somatic tissues as well as in the germline. We showed that general somatic tissue depletion led to the strict L1 arrest phenotype identical to that seen in *asna-1* mutants depleted from both maternal and zygotic contributions. We also found that intestine-specific depletion of ASNA-1 led to a growth defect but that it was not as severe as pan-somatic depletion, raising the possibility that the contribution of ASNA-1 to the L1 arrest likely came from both neurons and the intestine. This was further confirmed in double mutant by simultaneous depletion of ASNA-1 in intestine and neurons. Pan-somatic depletion after embryogenesis showed that *asna-1* was required for the development of worms from the L1 to the L2 stage but not from the L4 larval stage to adulthood. This indicated that ASNA-1 cannot be regarded as a housekeeping gene whose function is needed in a constitutive manner. Further, the analysis following auxin-mediated degradation showed that ASNA-1 function in germline was not required for the development and growth of worms but did have some effect on fertility. The relatively mild effect on fertility when ASNA-1 was depleted from the L1 stage onwards might point to the role of somatic gonadal tissues in this process. Alternatively, the failure to see a strong fertility defect might be due to the germline promoter used for the depletion analysis (Chen et al., 2020). Our earlier work showed that the insertion of SEC-61β to the ER membrane by ASNA-1 was critical for response to cisplatin. It is likely that when we deplete ASNA-1 from the L4 stage onwards sufficient SEC-61β has already been inserted into the ER membrane so that any further depletion will not affect SEC-61β levels and thus have no effect on cisplatin sensitivity. Depleting ASNA-1 earlier would prevent worms from reaching adulthood and not allow us to perform the adult stage cisplatin sensitivity.

The million mutation project (MMP) is a powerful resource for exploring mutations in almost every *C. elegans* gene (Thompson et al., 2013). It led to a collection of a large number of mutations in over 2000 whole genome sequenced strains. We tested seven single-point mutants in ASNA-1 generated by this project for cisplatin sensitivity (Table S1). Since all these seven strains were homozygous for the point mutants and were fertile, it was very likely that none of the seven point mutants affected reproduction. Given that the mutants were fertile we expected only mild changes in the overall effect on the function of the mutant ASNA-1. However, we cannot rule out the possibility that a compensatory second site mutant might act in some cases. Only two strains of the seven strains were cisplatin sensitive. We concluded that heavy mutation load *per se*, which is present in all the strains will not result in cisplatin sensitivity, and worms that were unhealthy were not inherently prone to cisplatin sensitivity due to some non-specific reason. In the screen of viable ASNA-1 mutants, which would be able to separate the *asna-1* clinically relevant functions in insulin secretion and cisplatin resistance, we identified *asna-1(A63V)* as important in function separation. This mutant preferentially exists in the oxidized state, has a strong cisplatin sensitivity phenotype and tail-anchored protein insertion defect, and perturbed subcellular localization. Importantly, the mutation in this particular alanine did not affect *C. elegans* IIS activity nor caused any growth or developmental problems. This allowed us to genetically separate *asna-1* functions even further without compromising the viability of the worms. Evidence for functional separation emerged from the earlier study of another point mutant: *asna-1(ΔHis164)*. We had previously shown that this mutant too is cisplatin sensitive and the mutant protein preferentially accumulates in the oxidized state (Raj et al., 2021). Here we present evidence that the growth-promoting functions of ASNA-1 remain largely intact in this mutant and the germline development is substantially better than that seen in the *asna-1* deletion mutant since oocyte and sperm are formed and ovulation (the exit of oocytes from the ovary) also occurs. While it is true that signals from sperm are needed for initiating the process, it is known that immature spermatids can also induce ovulation (McCarter et al., 1999) and therefore it is possible to have proper ovulation but no fertilization due to defects in sperm development.

*In silico* modeling showed that the A63V mutation, which is close to the Switch I motif, affects the coordination of a conserved Threonine in the P-loop motif with possible effects on ATP binding and hydrolysis. Further, another function separating mutation in highly conserved Histidine 164 lies in the Switch II motif and has no effect on insulin secretion and signaling. This indicates that Switch I and Switch II and P-loop motifs are excellent druggable targets that increase cisplatin sensitivity while sparing the insulin secretion role of ASNA-1.

## ACKNOWLEDGEMENTS

We thank the Caenorhabditis Genetic Center (funded by NIH Office of Research Infrastructure Programs P40 OD010440) and National Bioresource Project for the Experimental Animal “Nematode C. elegans” for providing strains.

## AUTHOR CONTRIBUTIONS

D.R., A.P-F., and G.K designed and performed the experiments, D.R., A.P-F., P.G. and G.K analyzed the data. D.R. and G.K wrote the paper. All authors critically revised the manuscript for intellectual content and approved the final version.

## COMPETING INTERESTS

The authors declare no competing interests.

## FUNDING

The work was supported by grants from the Swedish Cancer Society CAN 2018/664 (P.N.) and ALF means nr: ALFGBG-722971 (P.N.); Stiftelsen Assar Gabrielssons Fond FB19-44 (DR) and Stiftelsen Assar Gabrielssons Fond FB20-32 (DR).

## DATA AND MATERIALS AVAILABILITY

All data is available in the main text or supplementary materials.

## MATERIAL AND METHODS

### *C. elegans* strains maintenance and synchronization

The Bristol strain (*N2*) was the wild-type. The *asna-1(gk592672)* containing strain VC40357 was outcrossed 13 times using *unc-32(e189)* and *oxTi719* as balancers and kept *in trans* to the *hT2(qIs48)* balancer. ASNA-1^A63V^::GFP was generated by mutagenesis of pVB222GK (Kao et al., 2007) to introduce the A63V change using the QuickChange II Kit (Agilent Technologies). The resulting plasmid (pGK200) was used to generate the extrachromosomal array *rawEx8*. Genomic integration of *rawEx8* yielded *rawIs16. rawIs16* worms were outcrossed 4 times before analysis. The single-copy *asna-1::gfp (knuSi184)* contained 1.4 kb upstream promoter sequence driving the genomic *asna-1* coding region fused to GFP just before the stop codon followed by the *tbb-2* 3’ UTR. This construct was inserted on chromosome II at the *ttTi5605* locus using MosSCI technology (Frøkjær-Jensen et al., 2008) by Knudra Transgenics. *rawEx64* expressed *vha-6p*-*3xFlag::SEC-61β::opsin* as an extrachromosomal array and was outcrossed 2 times before use (Raj et al., 2021). All *C. elegans* strains used in this study are listed in **Table S2**. Worms were cultured under standard conditions at 20ºC on nematode growth media (NGM) plates unless stated otherwise and the *E. coli* strain OP50 was used as a food source. *Synchronization:* Synchronous larval populations were obtained by gravity separation as described previously (Oláhová et al., 2008).

### TAP-targeting analysis

The assay was performed as described previously (Raj et al., 2021). In short, live 1-day old adult animals were sedated in 1 mM Levamisole/M9 and mounted onto 2% agarose pads. The int8 and int9 cells in the posterior intestine were imaged. The fluorescence signals were analyzed at 488 nm and 555 nm by the LSM700 Confocal Laser Scanning Microscope (Carl Zeiss) with LD C-Apochromat 40x/1.1 W Corr. objective. Image processing of Z-stacks was performed with the ZEN Lite program (Zeiss). Correlation quantification was done using Volocity software (PerkinElmer). Correlation quantification was done using the Automatic Thresholding (Costes et al., 2004) method to set thresholds objectively.

### Insulin assays

Worms harboring integrated *daf-16::gfp (zIs356)* and *daf-28::gfp (svIs69)* arrays were grown at 20°C and imaged using a Nikon microscope, equipped with Hammamatsu Orca flash 4.0 camera. *daf-16::gfp* animals were analyzed within 10 minutes after mounting to avoid artifacts due to stress. DAF-28::GFP uptake by coelomocytes was scored in adult worms as described previously (Kao et al., 2007).

### Glycosylation analysis of tagged SEC61β

Young adult worms expressing *3xFlag::SEC-61β::opsin* from the *rawEx64* transgene were homogenized and taken for protein concentration determination. All samples were solubilized in Laemmli buffer, separated by SDS-PAGE followed by western transfer and detection by immunoblotting with anti-Flag (F1804, Sigma) antibody.

### Western blot analysis

*Reducing SDS-PAGE:* Synchronized young adult worms were homogenized and protein concentration was determined using the BCA assay (Thermo Scientific). Samples were boiled for 10 min in reducing loading buffer (SDS/ β-mercaptoethanol). Proteins were separated by SDS-PAGE and blotted onto PVDF membranes. *Non-reducing SDS-PAGE:* Lysates were boiled for 10 min in non-reducing (without β-mercaptoethanol) loading buffer and cooled to room temperature for 10 min. To protect free cysteine thiols from post-lysis oxidation, iodoacetamide was added to the samples at a final concentration of 25 mM followed by a 30 min incubation in darkness at room temperature. Proteins were separated by SDS-PAGE and blotted onto nitrocellulose membranes. *Antibodies:* anti-ASNA-1 antibody (Kao et al., 2007), anti-GFP antibody (3H9, Chromotek), and anti-Flag M2 antibody (F1804, Sigma) were used. To assess equal loading, membranes were stripped and probed with anti-alpha tubulin (T5168, Sigma). Band quantification was performed using ImageJ software (Schneider et al., 2012).

### Subcellular fractionation

Young adult animals grown at 20°C were harvested, washed, and lysed in extraction buffer (50 mM Tris, pH7.2, 250 mM sucrose, 2 mM EDTA). Supernatants were centrifuged for 60 min at 100,000xg at 4°C. The supernatant fraction was concentrated using Vivaspin Concentrators (Sigma). The pellet fraction was resuspended in 1x Laemmli buffer (Biorad). Proteins in both fractions were separated by SDS-PAGE and blotted onto PVDF membranes. Proteins were detected using anti-GFP antibody (3H9, Chromotek) or anti-alpha tubulin (T5168, Sigma).

### RNA isolation and quantitative PCR

Total RNA was extracted using Aurum Total RNA Mini Kit (BioRad). cDNA was synthesized using iScript cDNA Synthesis Kit (BioRad). qPCR was performed on a CFX Connect machine (BioRad) instrument using KAPA SYBR FAST qPCR Kit (KapaBiosystems) with the comparative Ct method and normalization to the housekeeping gene *F44B9*.*5*. All samples were tested in triplicates.

### Auxin treatment

Animals were transferred to NGM plates containing indicated concentration of the water-soluble auxin derivative naphthaleneacetic acid (N610; Phyto-Tech Labs). The auxin-containing agar plates were prepared on the day of use from a freshly made 800mM stock solution in water and stored in the dark.

### Cisplatin sensitivity assay

Cisplatin plates were prepared as described previously (Raj et al., 2021). L4 larvae were isolated and grown for 24h before exposure to cisplatin. After 24h cisplatin exposure, death was determined by the absence of touch-provoked movement when stimulated by harsh touch using a platinum wire.

### ASNA-1 model prediction

Three models for the *C. elegans* ASNA-1 (CELE_Zk637.5 #1-342) were obtained using the 3D modeling prediction program MODELLER (Šali and Blundell, 1993), having as a template the structure of the Get3 protein complex of *Chaeromium thermophilum* binding AMPPNP a non-hydrolyzable ATP agonist (PDB:3IQW) (Bozkurt et al., 2009). Two of the models were obtained by loading the sequence with one of the following mutations: A63V and ΔH164. The third structure does not include any mutation and mimics the wild-type protein. The model was assembled as a dimer and in complex with AMPPNP, Mg and Zn mimicking the structure obtained for *Chaeromium thermophilum* (PDB:3IQW).

## Notes

### Competing Interest Statement

The authors have declared no competing interest.

